# Generation and characterization of a patient-specific human induced pluripotent stem cell line from a Skogholt syndrome patient (ASCFi003-A)

**DOI:** 10.64898/2026.08.29.747981

**Authors:** Weronika Przybala, Swapnil Gupta, Hege Brincker Fjerdingstad, Per Selnes, Kulbhushan Sharma

## Abstract

We report the generation and characterization of a human induced pluripotent stem cell (iPSC) line derived from dermal fibroblasts of a patient with Skogholt’s disease, a rare maternally inherited neurodegenerative syndrome associated with choroid plexus dysfunction and impaired cerebrospinal fluid (CSF) homeostasis. Patient fibroblasts were reprogrammed using the non-integrating Repro-OSKGM kit. The resulting iPSC line exhibited typical pluripotent morphology, expressed canonical pluripotency markers, maintained a normal karyotype, retained the disease-associated genetic variant, was mycoplasma-free, and demonstrated trilineage differentiation potential. We also made choroid plexus (ChP) like organoids from the generated iPSCs. This patient-specific iPSC line provides a valuable resource for generating choroid plexus organoids and neurons to investigate disease mechanisms and develop therapeutic strategies.

## Resource table

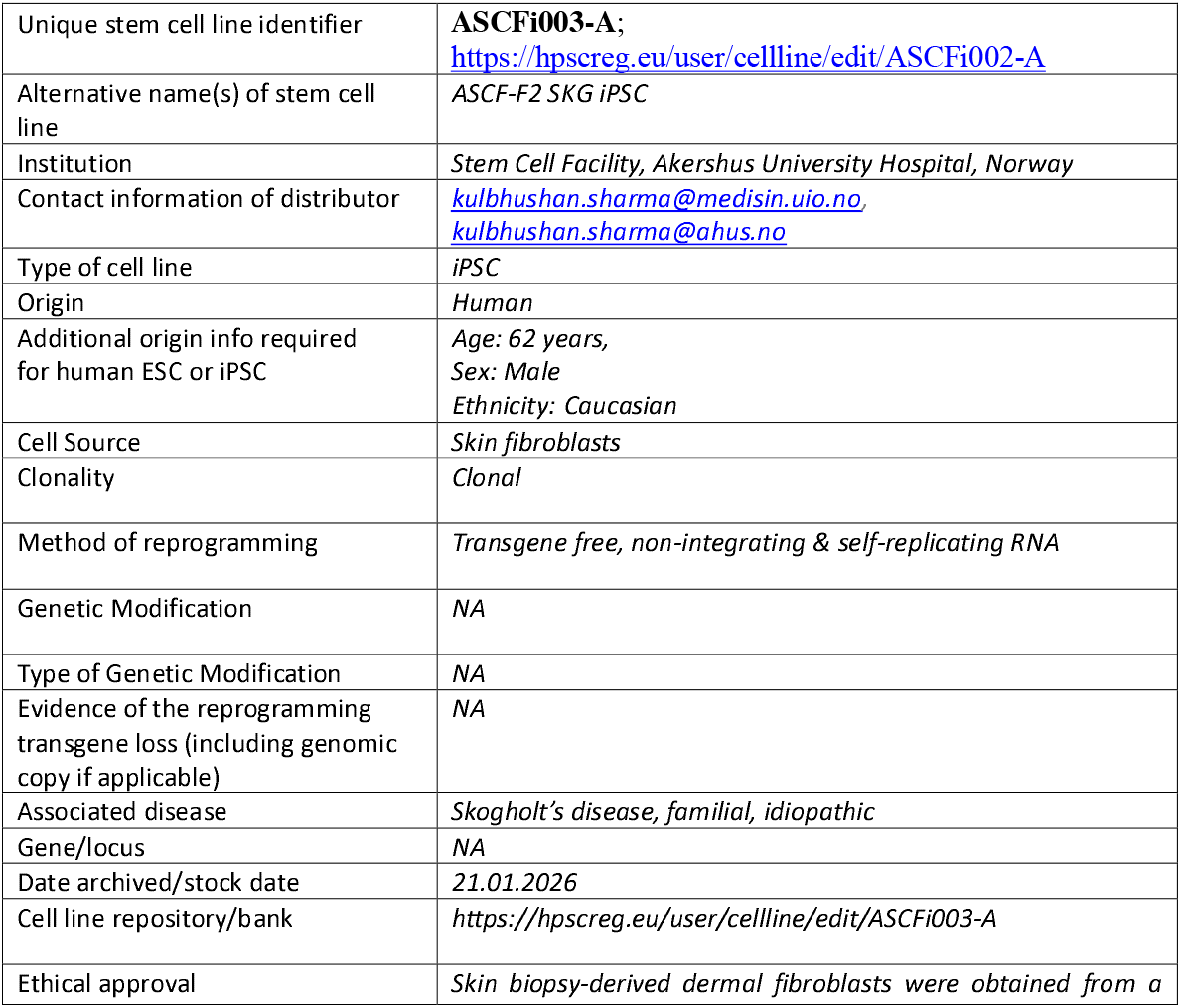
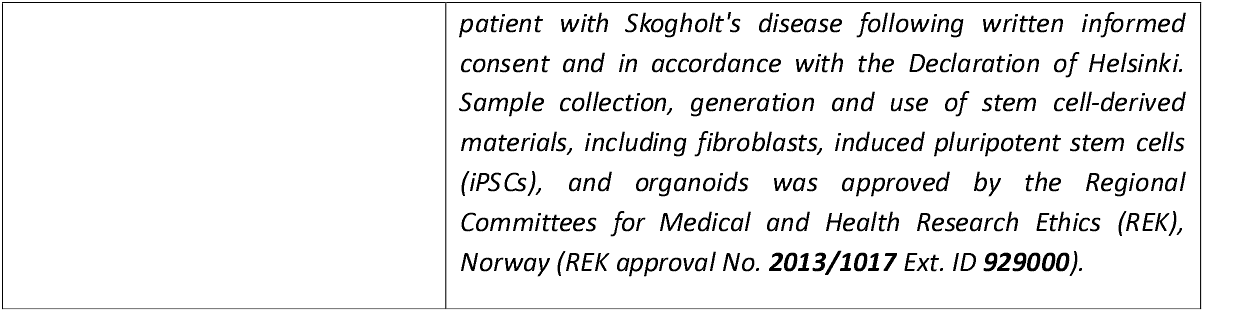

## Resource utility

This patient-derived iPSC line provides a unique human model of Skogholt’s disease, enabling generation of disease-relevant cell types, including choroid plexus epithelial cells, organoids, and neurons. The resource facilitates investigation of choroid plexus dysfunction, impaired CSF homeostasis, mitochondrial abnormalities, disease mechanisms, and potential therapeutic strategies.

## Resouce details

Skogholt’s disease is a rare maternally inherited neurodegenerative syndrome characterized by progressive neurological impairment **(1)**. Recent clinical, imaging, and biomarker studies indicate that the disease is associated with primary choroid plexus (ChP) dysfunction, resulting in reduced cerebrospinal fluid (CSF) production, impaired blood–CSF barrier transport, and defective CSF clearance in the absence of overt cortical atrophy or blood–brain barrier disruption. These observations suggest that impaired CSF homeostasis may represent an early pathogenic mechanism underlying the disorder **(1)**.

Importantly, a recent multimodal clinical study of individuals with Skogholt’s disease provided convergent in vivo evidence for a choroid plexus-centred clearance defect, further supporting a primary role of ChP dysfunction in the disease **(2)**. Affected individuals showed markedly reduced ChP volume and prolonged contrast washout on dynamic contrast-enhanced MRI, consistent with impaired ChP clearance dynamics, while no evidence of cerebral parenchymal blood–brain barrier disruption or overt cortical atrophy was observed. In addition, CSF concentrations of several neurological biomarkers, including tau, amyloid-β species, β-trace protein, neurofilament light chain, and glial fibrillary acidic protein, were substantially elevated despite normal or reduced plasma levels, indicating a pronounced disruption of CSF–plasma coupling and impaired clearance of solutes from the CSF compartment. The study also identified changes in trace-metal profiles suggestive of impaired ATP-dependent transport and reported a rare homoplasmic mitochondrial MT-RNR2 (m.1681G>A) variant that segregated with disease, providing further support for a potential mitochondrial contribution to ChP dysfunction. Collectively, these findings suggest that defective ChP transport and CSF clearance may constitute a primary disease mechanism in Skogholt’s disease and provide a strong rationale for investigating patient-derived ChP models as a means of defining the cellular and molecular consequences of the disease-associated mitochondrial defect **(2)**.

To establish a human in vitro model of Skogholt’s disease, dermal fibroblasts obtained from a skin biopsy of a clinically diagnosed 62-year-old male patient were reprogrammed into induced pluripotent stem cells (iPSCs) using the self-replicating RNA-based ReproRNA-OKSGM Kit (STEMCELL Technologies). Individual iPSC colonies were subsequently isolated and expanded as independent clones **(Table 1)**. Emerging colonies displayed characteristic human pluripotent stem cell morphology, including compact colonies with well-defined borders, a high nucleus-to-cytoplasm ratio, and prominent nucleoli.

**Table 1:**
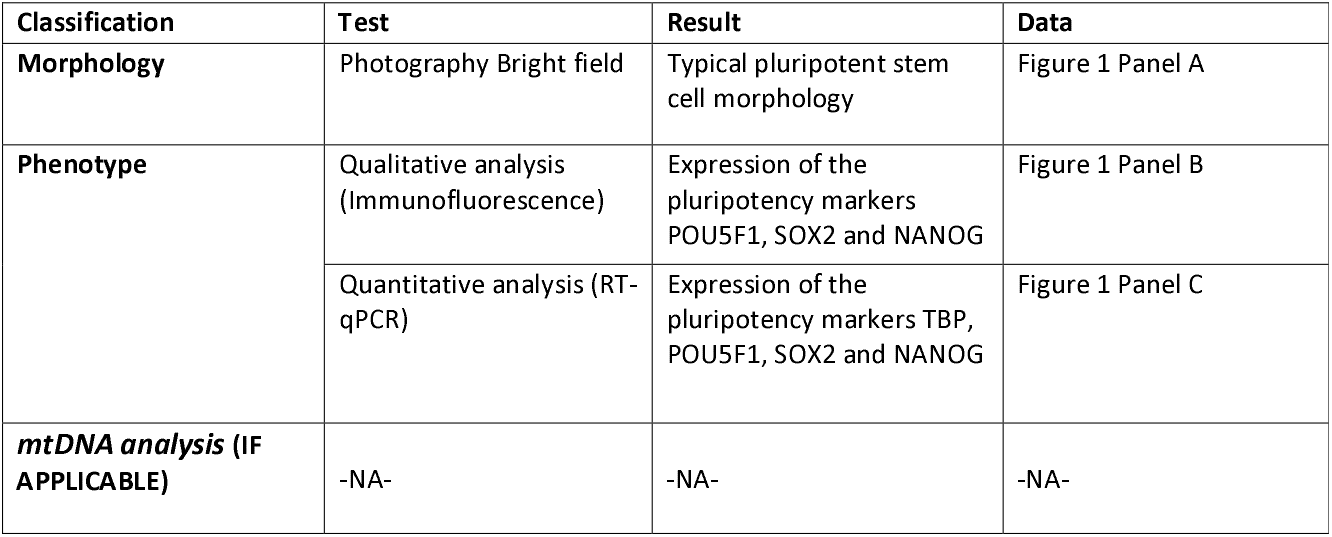
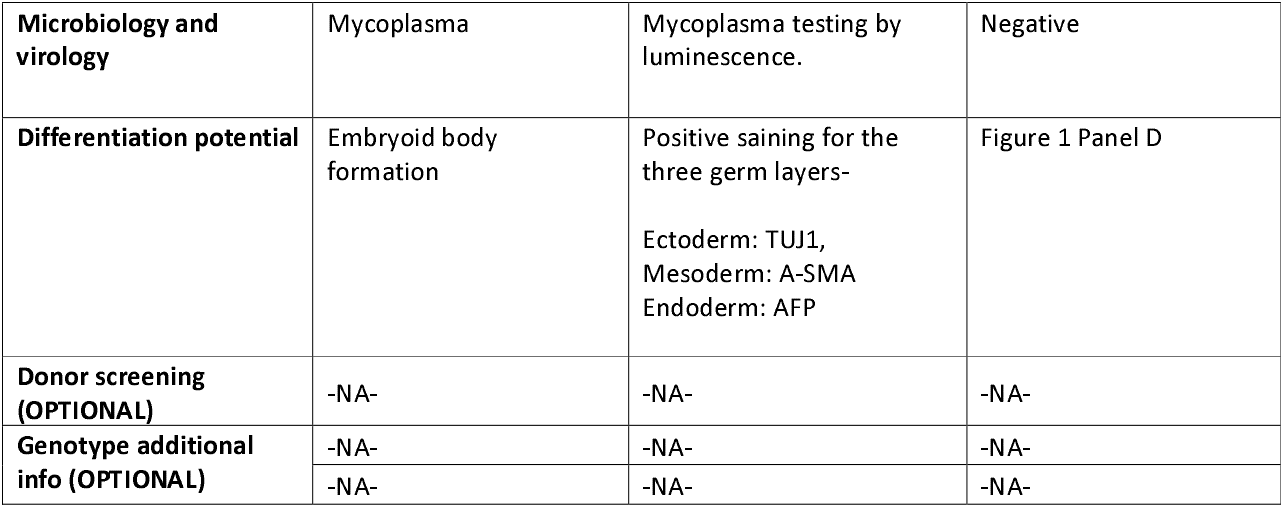
Characterization and validation.

| Classification | Test | Result | Data |
| --- | --- | --- | --- |
| **Morphology** | Photography Bright field | Typical pluripotent stem cell morphology | Figure 1 Panel A |
| **Phenotype** | Qualitative analysis (Immunofluorescence) | Expression of the pluripotency markers POU5F1, SOX2 and NANOG | Figure 1 Panel B |
|  | Quantitative analysis (RT-qPCR) | Expression of the pluripotency markers TBP, POU5F1, SOX2 and NANOG | Figure 1 Panel C |
| **mtDNA analysis (IF APPLICABLE)** | -NA- | -NA- | -NA- |
| **Microbiology and virology** | Mycoplasma | Mycoplasma testing by luminescence. | Negative |
| **Differentiation potential** | Embryoid body formation | Positive staining for the three germ layers-Ectoderm: TUJ1,<br>Mesoderm: A-SMA<br>Endoderm: AFP | Figure 1 Panel D |
| **Donor screening (OPTIONAL)** | -NA- | -NA- | -NA- |
| **Genotype additional info (OPTIONAL)** | -NA- | -NA- | -NA- |
|  | -NA- | -NA- | -NA- |

The established iPSC line, **ASCFi003-A**, exhibited typical pluripotent stem cell morphology (**Fig. 1A**; scale bar, 100 μm) and expressed the pluripotency markers POU5F1 (OCT4), SOX2, and NANOG, as confirmed by immunofluorescence (**Fig. 1B**; scale bar, 100 μm). POU5F1 and SOX2 showed the expected nuclear localization, supporting maintenance of the pluripotent state. Pluripotency was further assessed by RT-qPCR, which demonstrated expression of POU5F1, SOX2, and NANOG comparable to that of a positive-control iPSC line, using TBP as the reference gene (**Fig. 1C**). Functional pluripotency was confirmed by directed trilineage differentiation. Immunofluorescence analysis demonstrated differentiation into derivatives of all three embryonic germ layers, including TUJ1 (ectoderm), α-SMA (mesoderm), and AFP (endoderm) (**Fig. 1D**, scale bar, 100 μm). RTqPCR analysis was performed to detect the majority of recurrent karyotypic abnormalities reported in human embryonic stem cells (hESCs) and induced pluripotent stem cells (iPSCs) using the hPSC Genetic Analysis Kit (STEMCELL Technologies), according to the manufacturer’s instructions (**Fig. 1E (3)**). Routine mycoplasma testing confirmed that ASCFi003-A was mycoplasma-free, while sterility testing verified the absence of bacterial and fungal contamination (data not shown).

**Figure 1:**
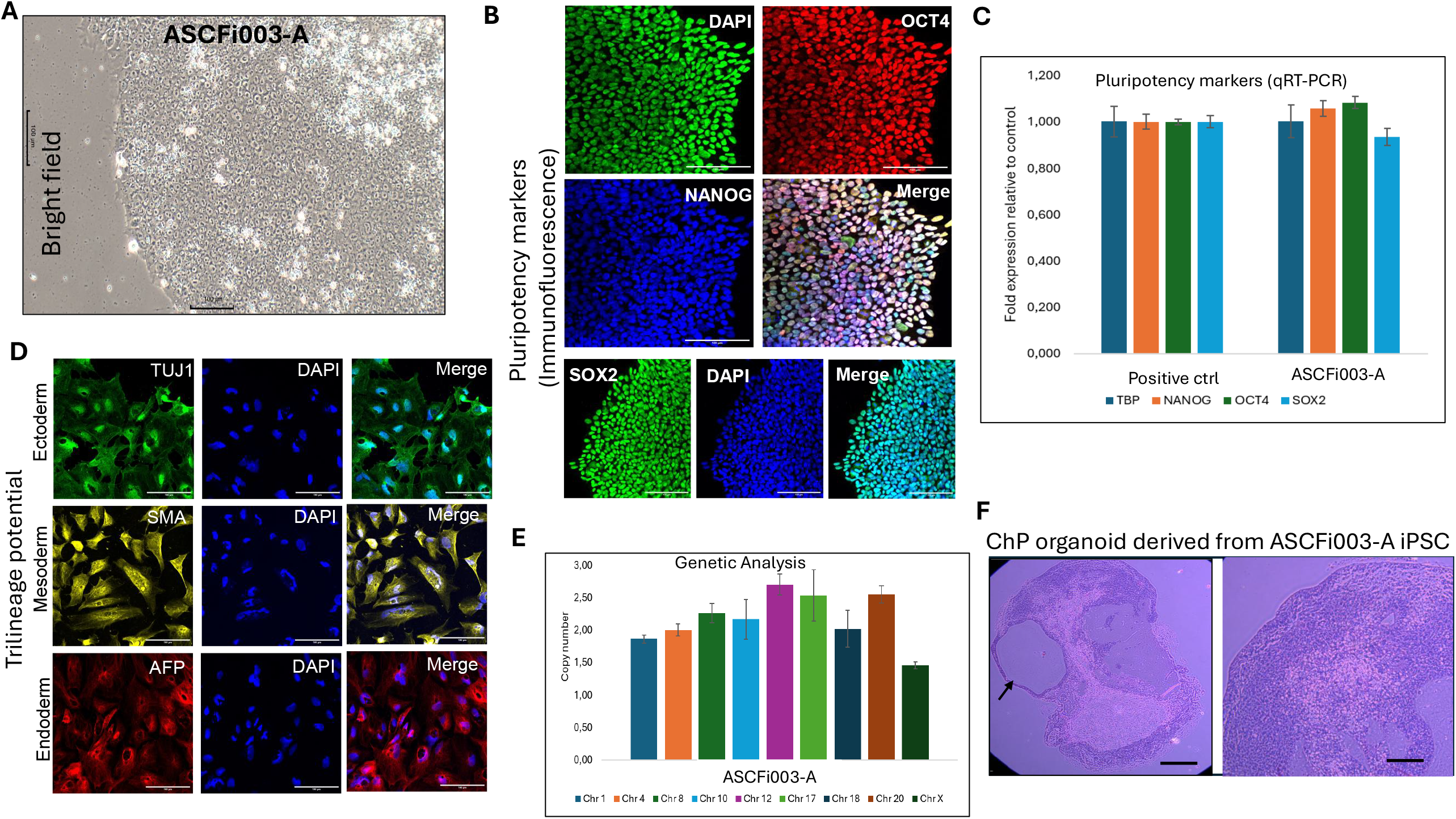
Characterization of the ASCFi003-A Skogholt’s disease iPSC line. **(A)** Representative bright-field image showing the morphology of the generated ASCFi003-A iPSC colonies. Scale bar: 300 µm. **(B)** Immunofluorescence staining demonstrating expression of the pluripotency markers OCT4, NANOG, and SOX2. Scale bar: 300 µm. **(C)** RT-qPCR analysis showing expression of the pluripotency-associated genes OCT4, NANOG, and SOX2. **(D)** Assessment of the trilineage differentiation potential of the ASCFi003-A iPSC line, demonstrating its capacity to differentiate into derivatives of the three embryonic germ layers. Scale bar: 300 µm. **(E)** Genetic characterization of the ASCFi003-A iPSC line. **(F)** Generation of choroid plexus (ChP) organoids from the ASCFi003-A iPSC line. Thirty-five-day-old ChP organoids were sectioned and stained with hematoxylin and eosin (H&E). The left image shows a representative section of the complete ChP organoid displaying a prominent cavity-like structure. Scale bar: 900 µm. The right image shows a higher-magnification view highlighting the characteristic epithelial morphology and organization of the ChP organoid. Scale bar: 900 µm.

The patient-derived iPSC line ASCFi003-A represents a valuable resource for investigating the cellular and molecular mechanisms underlying Skogholt’s disease. The line can be differentiated into disease-relevant cell types, including choroid plexus epithelial cells, neurons, glial cells, and brain organoids, providing opportunities to investigate processes associated with CSF production, blood–CSF barrier function, mitochondrial dysfunction, and clearance-dependent biomarker biology. To assess the potential of ASCFi003-A to generate choroid plexus (ChP) organoids, we followed the protocol described by Pellegrini et al (**4**). Following 30 days of differentiation, organoids were processed for hematoxylin and eosin (H&E) staining. Histological analysis revealed the formation of distinct internal cavities within the organoids, consistent with the development of fluid-filled luminal structures (**Fig. 1G**, scale bar, 300 μm); however, further characterization is required to determine their cellular identity and composition. Additional studies, including molecular and immunofluorescence characterization of ChP-specific markers and assessment of CSF-related functions, are ongoing. Furthermore, ASCFi003-A provides a platform for generating CRISPR-based isogenic controls, performing mechanistic studies, identifying disease-associated biomarkers, and evaluating potential therapeutic strategies targeting choroid plexus dysfunction and impaired CSF clearance.

## 3. Materials and Methods

### 3.1. Reprogramming skin fibroblasts into iPSCs using ReproRNA-OKSGM

Skogholt patient biopsy was obtained at the Ahus stem cell facility. The biopsy was washed and processed for fibrblast growth in DMEM/F12 media (Gibco) with 10% FBS (Gibco). After 2 weeks of growth, the fibroblasts were passaged. Passage 3 fibroblasts were used for reprogramming. Reprogramming was done using intregration-free ReproRNA-OKSGM Kit (STEMCELL Technologies) according to the manufacturer’s protocol. The iPSC colonies appeared after three weeks. These iPSC colonies were manually transferred to Matrigel-coated dish for isolation and expansion, and were maintained in mTeSR media until stable (STEMCELL Technologies) at 37⍰C and 5% CO2. **Table 1** summarizes the morphological characteristics of the derived iPSC colonies along with the results of their molecular and cellular characterization.

### 3.2. Immunofluorescence staining

The iPSCs at passage 10 were fixed in 4% paraformaldehyde (PFA) for 6 min at room temperature. Following three washes with phosphate-buffered saline (PBS), the cells were blocked and permeabilized using 1 ×PBS containing 5% bovine serum albumin (BSA) and 0.4% Triton X-100 for 1 h at room temperature. Primary antibodies were incubated overnight at 4⍰C (detailed in **Table 2**). The following day, after three washes with PBS (5 min per wash), a secondary antibody (Goat Anti-Rabbit AF488) was incubated for 1 h at room temperature in the dark. After three additional PBS washes (5 min per wash), nuclei were counterstained with DAPI for 7 min at room temperature in the dark, followed by a final PBS rinse. Images were acquired using LSM 500 (Carl Zeiss) and ZEN black software.

**Table 2:** Reagents and equipment used for characterization of pluripotency and embryoid body differentiation.

| **Characterization** | **Target / reagent** | **Host / type** | **Dilution** | **Manufacturer** | **Catalog No.** | **RRID** |
| --- | --- | --- | --- | --- | --- | --- |
| **ICC – Pluripotency** | *OCT4* | *Rabbit* | *1:100* | *Stemgent* | *09-0023* | *AB_2167689* |
|  | *SOX2* | *Rabbit* | *1:100* | *Stemgent* | *09-0024* | *AB_2195775* |
|  | *NANOG* | *Mouse* | *1:400* | *Sigma-Aldrich* | *MABD24* | *AB_11203826* |
|  | *DAPI mounting medium* | --- | — | *Sigma-Aldrich* | *F6057-20ML* |  |
|  | *Cy2-conjugated secondary antibody* | *Goat* | *1:200* | *Jackson ImmunoResearch / AffiniPure* | *101-226-003* | *AB_2337310* |
|  | *Cy3/C3-conjugated secondary antibody* | *Goat* | *1:400* | *Jackson ImmunoResearch / AffiniPure* | *111-165-003* | *AB_2338000* |
| **ICC – Embryoid bodies** | *βIII-tubulin (TUJ1), ectoderm marker* | *Mouse* | *1:600* | *Sigma-Aldrich* | *T8578* | *AB_1841228* |
|  | *Alpha-fetoprotein (AFP), endoderm marker* | *Rabbit* | *1:100* | *Thermo Fisher Scientific* | *PA5-21004* | *AB_11157055* |
|  | *α-Smooth* *muscle actin (SMA), mesoderm marker* | *Goat* | *1:200* | *Novus Biologicals* | *NB300-978* | *AB_2273628* |
|  | *Cy2-conjugated secondary antibody* | *Goat* | *1:200* | *AffiniPure* | *101-226-003* | *AB_2337310* |
|  | *Cy3/C3-conjugated secondary antibody* | *Goat* | *1:400* | *AffiniPure* | *111-165-003* | *AB_2338000* |
| **Imaging** | *Confocal microscope* | — | — | *Carl Zeiss* | *LSM700* |  |
|  | *Imaging software* | — | — | *Carl Zeiss* | *ZEN Black, 2011* |  |
| **qPCR – Pluripotency** | **Target** |  |  | **Company** | **Cat no** | **Amplicon size** |
|  | *SOX2 TaqMan Gene Expression Assay* | — | — | *Thermo Fisher Scientific* | *4331182; Hs01053049_s1* | *91bp* |
|  | *NANOG TaqMan Gene Expression Assay* | — | — | *Thermo Fisher Scientific* | *4331182; Hs04260366_g1* | *99bp* |
|  | *POU5F1 (OCT4) TaqMan Gene Expression Assay* | — | — | *Thermo Fisher Scientific* | *4331182; Hs00999634_gH* | *64bp* |
|  | *Human ACTB endogenous control* | — | — | *Thermo Fisher Scientific* | *4333762F* | *171bp* |
|  | *TaqMan Fast Advanced Master Mix* | — | — | *Thermo Fisher Scientific* | *4444964* | — |
|  | *qPCR instrument* | — | — | *Thermo Fisher Scientific* | *QuantStudio 3* | — |

### 3.3. Quantitative reverse transcription polymerase chain reaction (qRT-PCR)

Total RNA was prepared using the Trizol Reagent (Invitrogen) according to manufacturer’s instruction. cDNA synthesis was carried out using a Superscript kit (Invitrogen). qRT-PCR reactions were prepared using TaqMan Fast Advanced Master Mix (Applied Biosystems) and TaqMan Gene Expression Assays. The gene-specific TaqMan assay IDs used in this study are listed in **Table 2**. qPCR was performed using QuantStudio 3 (Thermofisher Scientific) according to the manufacturer’s instructions. All samples were analyzed in technical replicates, and no-template controls were included to monitor nonspecific amplification and contamination.

### 3.4. In vitro trilineage differentiation

Pluripotency of iPSC was confirmed using directed differentiation protocol into each of three germ layer which then assayed using immunostaining and qPCR analysis. Endoderm differentiation was induced by basal differentiation media containing 20 ng/mL Activin A (Peprotech), and 3 μM CHIR (Stemgent) for 7 days. Mesoderm was induced using basal differentiation media containing 2 ng/mL Activin A (Peprotech) and 40 ng/mL BMP4 (Peprotech) for 7 days. Ectoderm differentiation was induced by basal differentiation media containing 10 μM SB-431542 (TOCRIS) and 100 ng/mL Noggin (Peprotech) for 12 days. For basal differentiation media, 1X N2 supplement (Gibco), 1X B27 without retinoic acid (Gibco), 1X Glutamax (Gibco), and 200 μM Ascorbic acid were supplemented into DMEM/F12 (Gibco).

### 3.4. Testing differentiation potential to choroid plexus (ChP) organoids

To assess the differentiation potential of the Skogholt patient-derived iPSC lines, we generated choroid plexus (ChP) organoids following the protocol described by Pellegrini et al. (**4, 5**). Briefly, iPSCs were dissociated and aggregated to generate three-dimensional embryoid bodies, which were subsequently subjected to a defined sequence of differentiation conditions to promote ChP epithelial specification. ChP fate was induced through modulation of developmental signaling pathways, including activation of WNT signaling and treatment with BMP4, which promote the acquisition of ChP-like epithelial characteristics. Following the initial patterning phase, the developing aggregates were transferred to maturation conditions and maintained in culture to allow further differentiation and organization of the ChP-like epithelium. During culture, organoid development was monitored by assessing changes in morphology, including the appearance of epithelial structures and characteristic cystic regions containing fluid-like material. The resulting organoids were subsequently evaluated for their morphological characteristics as an indication of successful ChP differentiation. This differentiation approach has previously been demonstrated to generate human ChP organoids that recapitulate key morphological and functional features of the native ChP, including the development of a polarized, barrier-forming epithelium and secretion of CSF-like fluid (**4, 5**).

## Acknowledgements

This work was supported by Akershus University Hospital (AHUS) internal funding for the stem cell unit.

